# Guanosine monophosphate reductase 1 is a potential therapeutic target for Alzheimer’s disease

**DOI:** 10.1101/215947

**Authors:** Hongde Liu, Kun Luo

## Abstract

Alzheimer’s disease (AD) is a severe neurodegenerative disorder. Identification of differentially expressed genes in AD would help to find biomarker and therapeutic target. Here, we carried out an analysis to identify the age-independent and AD-specific genes. We found that genes MET, WIF1 and NPTX2 are down regulated in AD. WIF1 and MET are in signaling of WNT and MET, regulating the activity of GSK3β, thus in AD. Importantly, we found gene GMPR shows a gradual increase in AD progress. A logistic model based on GMPR exhibits a good capacity in classifying AD cases. GMPR’s product GMPR1 links with AMPK and adenosine receptor pathways, thus associating phosphorylation of Tau in AD. This allows GMPR1 to be a therapeutic target. Therefore, we screened five possible inhibitors to GMPR1 by docking GMPR1 with 1174 approved drugs. Among them, lumacaftor is ideal due to its high affinity and light molecular weight. We then tested the effect of lumacaftor on AD model mice. After twenty days of oral administration, β-Amyloid accumulation is slowed down and phosphorylation of Tau is almost eliminated in the treated mice, showing a satisfying effect. In conclusion, the elevated expression level of GMPR tightly associates with AD progress and leads to AD phenotype probably through AMPK and adenosine receptor pathways; and one of therapeutic strategies is to inhibit GMPR’s product with lumacaftor.

**Significance Statement:** We found the elevated expression level of GMPR tightly associates with AD progress and leads to AD phenotype probably through AMPK and adenosine receptor pathways; and the therapeutic strategy targeting GMPR1 with lumacaftor shows a satisfying result.

## Introduction

Alzheimer’s disease (AD), the most common cause of dementia, is characterized by the extracellular amyloid plaques and the intraneuronal neurofilament tangles (NFT), being composed of beta-amyloid protein (Aβ) and phosphorylated Tau protein, respectively [1]. AD presents a complicated pathological mechanism, associating multiple pathways including Wnt signaling pathway and 5’ adenosine monophosphate-activated protein kinase (AMPK)-signaling pathway [2, 3].

Glycogen synthase kinase 3 (GSK3β) is one of the components of Wnt signaling and seemingly displays a central role in AD [3, 4]. Activation of Wnt signaling inhibits GSK-3β-mediated hyperphosphorylation of Tau protein, thus preventing the formation of NFT [3, 5]. Reversely, evidences also suggested that Aβ exposure induced GSK3β activity [6]. MET signaling regulates cell growth and development, thus in cancer. It has a crosstalk with Wnt signaling through β-catenin and GSK3β. MET makes a contribution to nuclear translocation of β-catenin by its tyrosine phosphorylation (by SRC) or by inhibition of GSK3β [7, 8]. The nuclear translocation of β-catenin results in transcriptional activation of Wnt ligand WNT7B and MET [3, 5].

AMPK sensors AMP/ATP ratio (ATP level) regulates the cellular energy metabolism. It is possible that the AMPK activity can decrease Aβ generation either through regulating neuronal cholesterol and sphingomyelin levels or through upregulation of BACE1, one kind of the enzymes that cleaves amyloid precursor protein (APP) [9, 10]. AMPK is also implicated in hyperphosphorylation of Tau protein [11]. In another pathway, the extracellular adenosine, which is generated from AMP through ecto-50-nucleotidase (CD73), binds to A1/2 receptor thus leading to ERK-dependent increase of Tau phosphorylation and translocation towards the cytoskeleton [12-14].

Identification of gene expression changes in AD will help to find molecular mechanisms of AD and to discover new drug targets [3]. The signaling pathways of Wnt, AMPK, MET and A1/2 enrich the expression-altered genes in AD, for instance, the decreased β-catenin [15], the elevated Dkk1 [16], the increased A1 and A2 receptors [13], the elevated AMP deaminase and the upregulated GSK3β [4, 17]. In an analysis on AD brains of postmortem human, Yusaku et al’s identified a number of downregulated genes including NPTX2 [18]. Xiao at al confirmed the reduction of NPTX2 in AD and suggested a mechanism that the reduced NPTX2 is probably due to an increase of miR-1271[19].

In the present study, we focused on two issues. One was to identify genes with a different expression in AD rather than in old population. It is accepted that AD is a kind of neurodegenerative disorder of old-aged humans. But in some of the old people, AD is not found even in those people who have a comparable age with the AD patients [20]. It is necessary to discriminate the age-dependent and age-independent in AD expression analysis, which will help to find new markers for AD. The other is to find new therapeutic targets. Current therapeutic targets are either to enhance neurotransmitter systems or to modify the pathways that cause disease [2]. The latter focuses on both Aβ and NFT by targeting secretase, neutral endopeptidase, endothelin-converting enzyme, vaccination, apolipoprotein E (ApoE), GSK3β, CDK5 and so on [21, 22].

Here, we carried out a comparative analysis to identify the genes that express differentially in AD. Gene GMPR, which encodes human guanosine monophosphate reductase 1 (GMPR1), was found gradual increase with AD progress. Targeting GMPR1, we discovered five possible inhibitors by docking GMPR1 with the Food and Drug Administration (FDA) approved drugs. For one of the inhibitors, lumacaftor, we evaluated the inhibiting effect in AD mice. Phosphorylation of Tau was almost eliminated in the treated AD mice.

## Results

### Identification of the age-independent differentially expressed genes

In dataset GSE36980, which includes 32 AD and 47 non-AD samples, we identified six down-regulated genes and one up-regulated gene under the criteria of both p-value ≤ 10^−5^ and the absolute value of log2 (fold change) ≥ 0.1 (Fig. 1A). In AD samples, the expression of genes NPTX2, WIF1, MET, LINC00643, CBLN4, CRHBP and PPEF1 are down regulated. The down regulation of NPTX2 and MET were previously reported in literatures [18, 19]. Gene GMPR, which encodes protein GMPR1, is up regulated in AD cases (Fig. 1A). The genes SYT5 and CHRNB2 have a significant p-value but a less fold change, thus they are not considered in further analysis.

**Figure 1.**
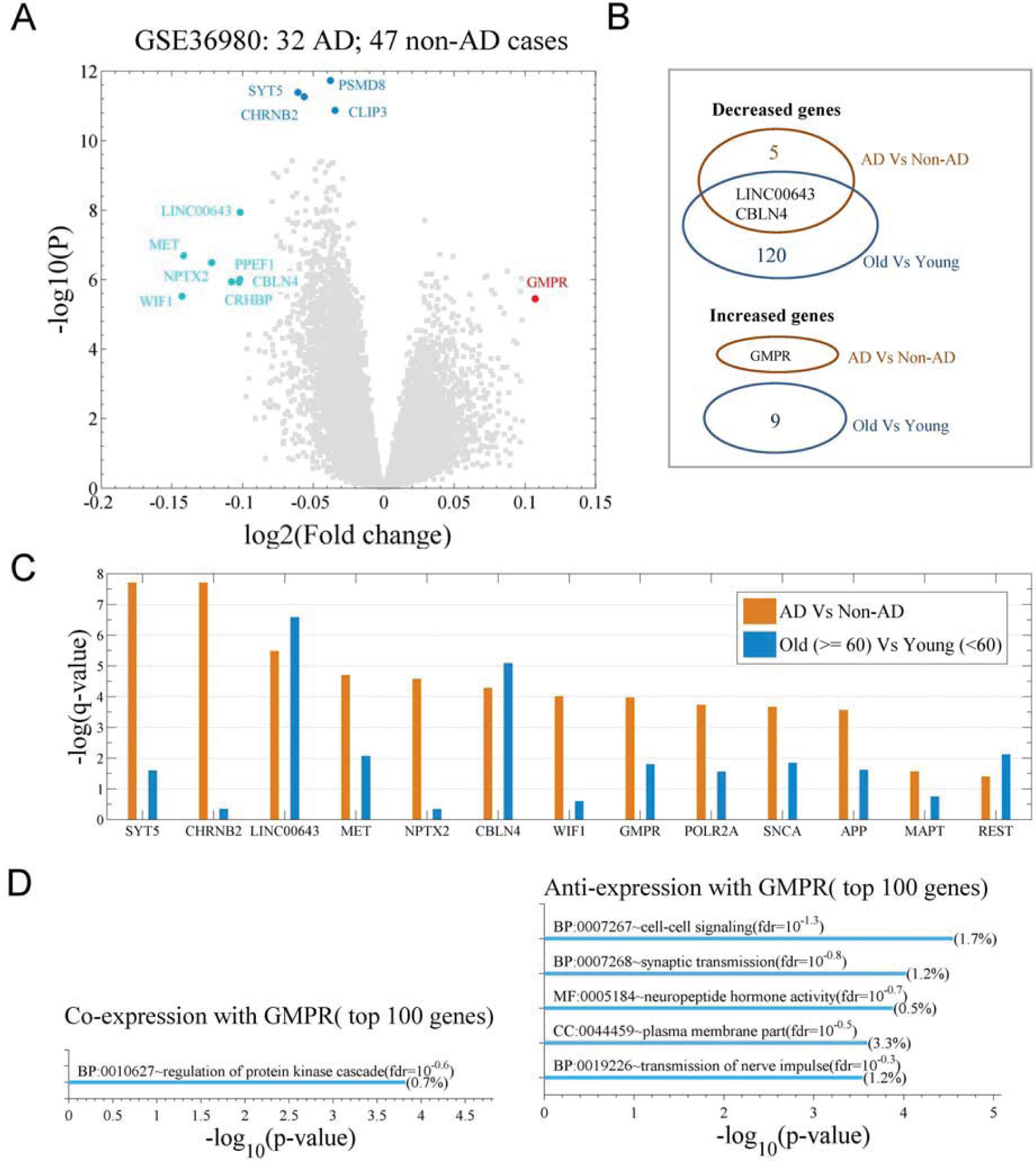
Differential expression analysis for the postmortem human brain tissues of Alzheimer’s disease (AD). The data used is the microarray data of 33297 human transcripts in 32 AD samples and 47 non-AD samples (GSE36980). **A**. A volcano plot (fold change vs. p-value (two-sample *t*-test)) for the AD vs. the non-AD cases, the cutoff for the differentially expressed genes is p-value ≤ 10–5 and either log_2_ (fold change) ≥ 0.1 or ≤ –0.1. **B**.Venn plot indicates the overlapped number of the genes with a different expression either in AD cases or in old population (Figure S1A). **C**. Shown are q-values of genes MAPT, APP, SNCA, POLR2A, GMPR, WIF1, NPTX2, MET, LINC00643, REST, SYT5 and CHRNB2 in two comparisons (AD vs. non-AD and old vs. young). **D**. An enrichment analysis for both top 100 co-expression genes (top panel) and top 100 anti-expression genes (bottom panel) of GMPR. The analysis is on the terms of gene ontology (GO) with DAVID. The co- and anti-expression genes are identified with the Pearson correlation coefficients (r) between GMPR and each gene on dataset GSE36980.

Between old and young population (GDS5204), genes LINC00643 and CBLN4 exhibit a different expression (Fig. 1SA and Fig. 1B-C), suggesting that the two are age-dependent in expression. We noticed that the fold change of the expression between AD and non-AD is not so great, indicating that AD is probably with chronic feature (Fig. 1SB). Interestingly, gene GMPR is up regulated in AD but not in the old population (Fig. 1C), suggesting that GMPR is an age-independent biomarker for AD. The enrichment analysis proved that the GMPR’s co- and anti-expression genes are in cell-cell signaling, synaptic transmission and regulation of protein kinase cascade (Fig. 1D), implicating an important role in the nerve cells.

### The models of classifying AD and non-AD cases

In order to evaluate the capacity of the differential expression genes being as biomarkers for AD, we constructed the logistic regression models with the genes and their combinations (Fig. 2A and Tab. S1-2). The models based on NPTX2, GMPR and MET exhibit a more than 0.8 of area under the curve (AUC) on dataset GSE36980 (32 non-AD and 47 AD cases) (Fig. 2B). The models with the genes combinations did not show an obvious enhancement in AUC (Fig. S2A). On dataset GSE28146 (8 non-AD and 22 AD cases), the models showed a weak capacity of classifying (Fig. S2B), which is due to a less difference between the control and the incipient cases.

**Figure 2.**
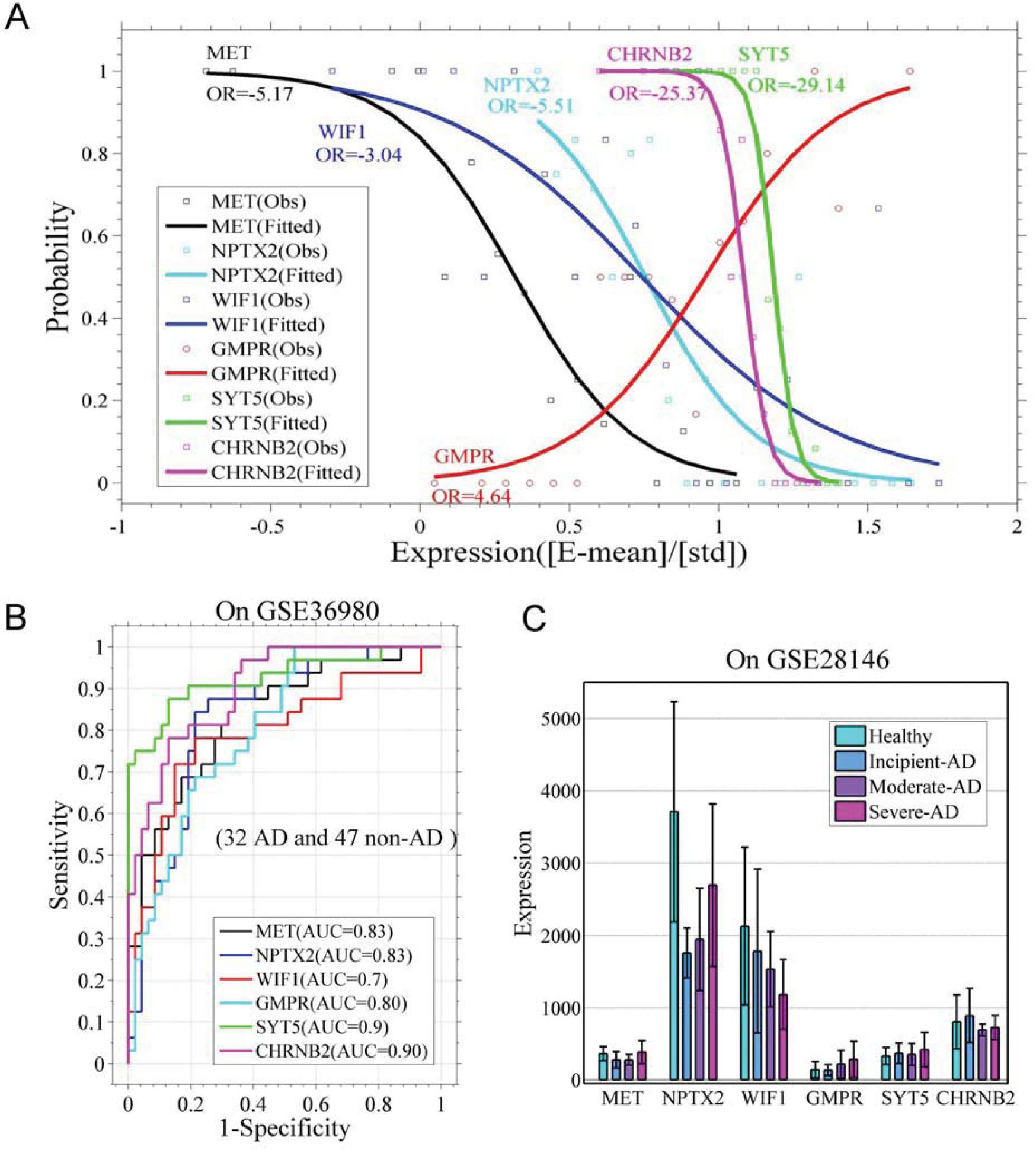
Performance of the logistic models based on the differentially expressed genes. **A**. Logistic regression models based on the differentially expressed genes GMPR, WIF1, NPTX2, MET, SYT5 and CHRNB2. Models’ parameters are listed in Table S1 and S2. **B**. The receiver operator characteristic (ROC) curves shows the performance of the models on dataset of GSE36980 (32 AD and 47 non-AD samples). AUC means area under the curve. **C**. Average expression level of the genes GMPR, WIF1, NPTX2, MET, SYT5 and CHRNB2 in 8 Non-AD, 6 slight-AD, 9 moderate-AD and 7 severe-AD cases (GSE28146), error bar shows the standard deviation.

Dataset GSE28146 is composed of 8 controls, and 7 incipient, 8 moderate and 7 severe AD cases, which allows us to observe the gene expression change with the progress of AD. On this dataset, we observed a gradual increase of GMPR and a gradual decrease of WIF1 from health to severe AD cases (Fig. 2C), demonstrating the two genes can be a well indicator for AD progress. Importantly, the increased expression of GMPR makes the product of GMPR (GMPR1) be a potential therapeutic target. Although the expression of NPTX2 is lower in AD cases than in control, it shows an increase from incipient to severe cases (Fig. 2C). MET, SYT5 and CHRNB2 do not show a distinct expression on this dataset (Fig. 2C).

Additionally, we investigated the transcription regulation for GMPR. At the enhancers of the gene, transcription factors (TFs) JUND and CBX3 have binding motifs (Fig. S3A). In the healthy cases GMPR and the TF show a negative correlation (r < –0.6) in gene expression; However, in the severe AD cases, they exhibit a positive correlation (r > 0.5) (Fig. S3B), indicating a reverse in transcriptional regulation in AD. Also, the expression of JUND is increased in AD (Fig. S3C). This means that the increased expression of GMPR is probably caused by an abnormal transcription regulation conducted by TF JUND.

Taken together, an elevated level of GMPR in AD was observed in both datasets. Gene GMPR and its product GMPR1 can be both a potential therapeutic target and a diagnosis biomarker.

### Links between the differentially expressed genes and AD pathomechanism

The genes with a differentia expression associate with AD amyloid plaques and NFTs through multiple paths. Protein MET is tyrosine kinase receptor that can be activated by binding of hepatocyte growth factor (HGF). MET signaling represses the activity of GSK3β, which is associated with the increase of phosphorylation of Tau [4]. The signaling of MET also contributes to the nuclear translocation of β-catenin, consequently promoting the transcription of genes WNT7 and MET (Fig. 3A) [4, 7, 8]. Down regulation of MET seems to facilitate AD development.

**Figure 3.**
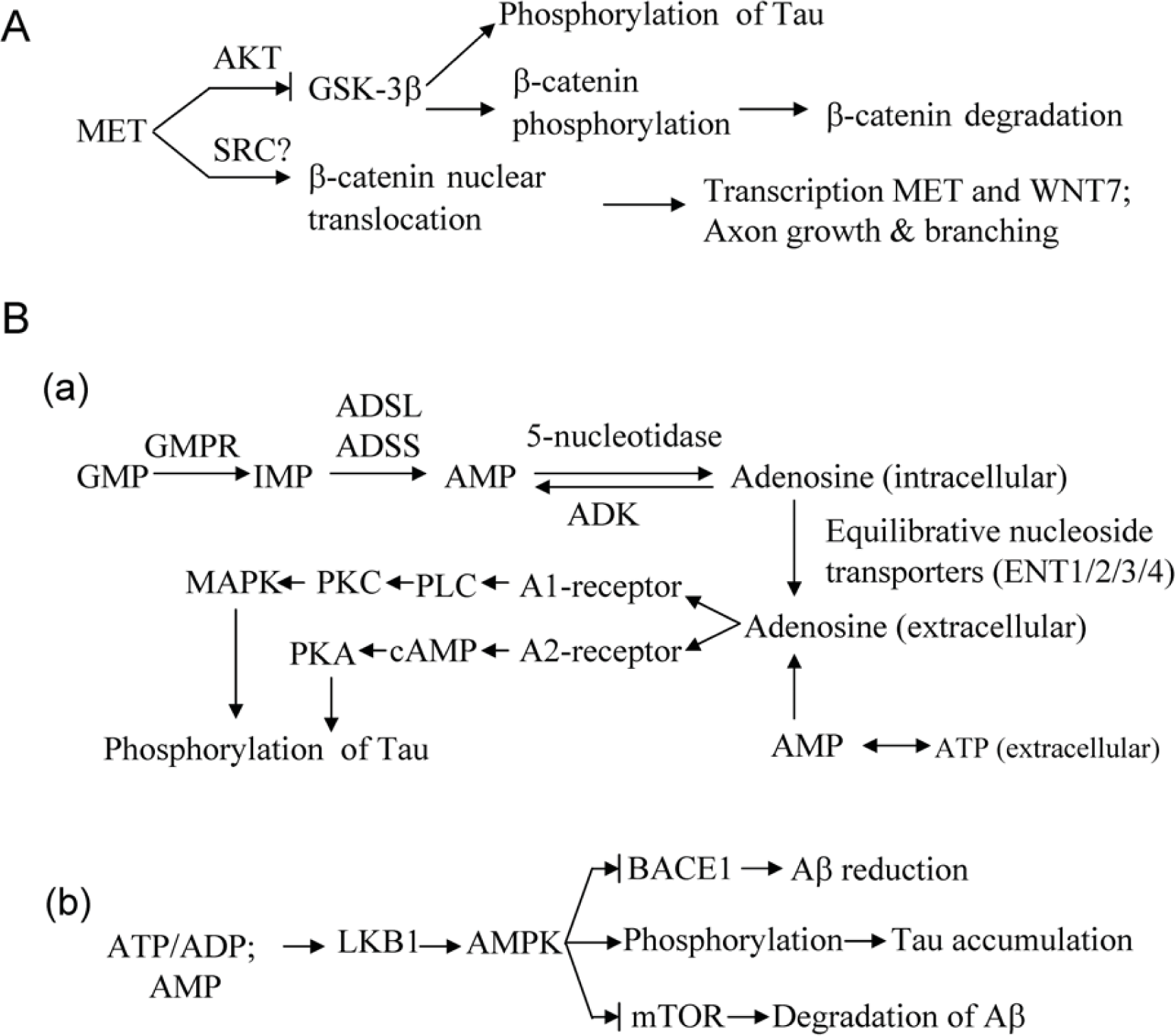
Possible mechanisms between the differentially expressed genes and Alzheimer’s disease. **A**. For MET. **B**. For GMPR.

Protein GMPR1, encoded by gene GMPR, is responsible for the conversion of GMP and NAPDH to IMP and ammonium, thus connecting purine metabolism, ATP generation and transcriptional signaling (Fig. S4A). There exist reversible conversions between IMP and AMP, between AMP into adenosine (A) [29]. There are two possible paths linking GMPR and AD phenotype. The first path is through the activations of A1/A2 receptors by extracellular adenosine [29]. Extracellular adenosine is either from transmembrane transport by equilibrative nucleoside transporters or from the conversion of extracellular AMP (Fig. 3B). Bindings of adenosine to A1/A2 receptors transduce the signaling to both PKA and PKC, thus leading the phosphorylation of Tau (Fig. 3B) [14]. In literature, an up-regulation of adenosine receptors was observed in the frontal cortex in AD [14]. Also, evidences suggested that noxious brain stimuli enhance the extracellular levels of adenosine [30]. It is speculated that an increased GMPR level will accumulate adenosine given that other enzymes remain no significant change. A model in which we let ATP concentration oscillate with time indicates that adenosine will increase with time (Fig. S4B). Also, AMP can be directly an agonist of A1 receptor [12]. The second possible path is through AMPK (Fig. 3B), which associates tau accumulation and influence Aβ generation [9, 11, 31]. Expression change of GMPR exerts an influence on the leves of IMP, AMP and ADP. It is well known that changes of ratio of ATP to ADP or to AMP can be sensed by serine/threonine-protein kinase LKB1, then activate AMPK pathway.

In short, the up-regulation of GMPR associates AD phenotype at least through two pathways, namely adenosine receptor-mediated pathway and AMPK pathway. That means the expression alteration of GMPR is probably a source more upstream and essential to AD. Treatment targeting GMPR or its product is to be a possible strategy for AD.

WIF1, which inhibits the Wnt signaling, is down regulated in AD, which leads an enhanced Wnt signaling, thus a low activity of GSK3β and a reduced phosphorylation of tau [5]. This means the down-regulation of WIF1 is a predictive response in neuron cells.

Regarding NPTX2, literature suggested that its reduction contributes to cognitive failure in AD through regulating GluA4 [19]. We still do not know the function of LINC00643, but it shows a higher expression in brain than in other tissue [32].

### The inhibitors to GMPR1

In order to explore a therapeutic strategy, we screened a possible inhibitor for enzyme GMPR1 from 1174 FDA-approved drugs. We hoped to repurpose one of the drugs to treat the AD. The 1174 drugs show different affinity in binding to GMPR1 (Fig. S5A). The drugs were ranked by the docking affinity (Fig. S5A; Tab. S3). By considering the molecule weight and the octanol-water partition coefficients (logP), five drugs can be candidates. Fig. 4 shows the full view and the interacting details for the five docked complex. Four drugs involve hydrogen-bond interaction (H-bond). Lumacaftor, which is able to correct the folding of cystic fibrosis transmembrane conductance regulator protein (CFTR) with F508del mutation [33], is ideal due to its small molecular weight (452), small topological polar surface area (97.8 Å2) and reasonable logP (4.37) (Tab. S3). Lumacaftor also shows a well docking with GMPR2, the paralog of GMPR1 (Fig. S5B), probably indicating that it can inhibit both GMPR1 and GMPR2.

**Figure 4.**
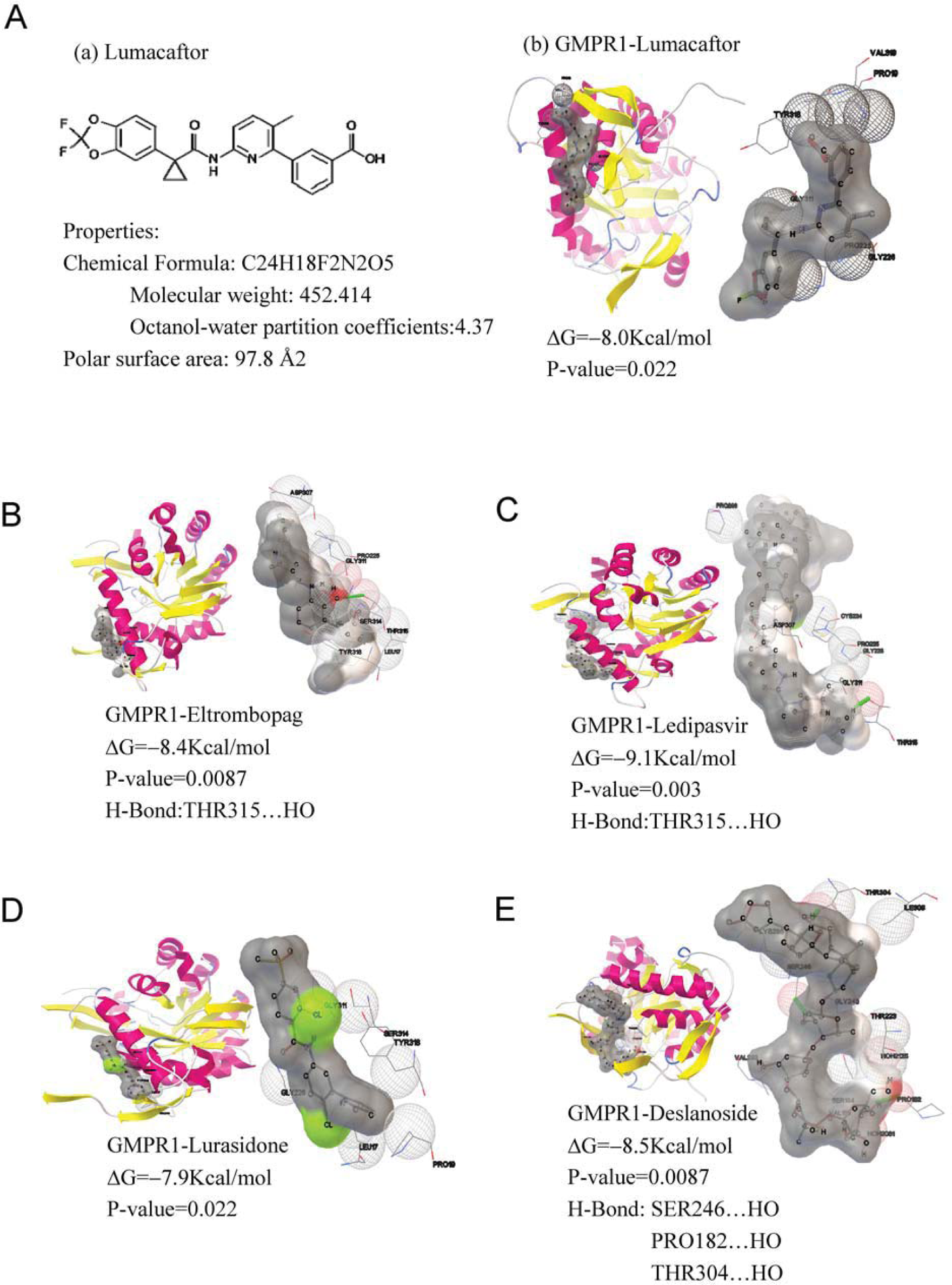
Five optimal drugs filtered by docking with human guanosine monophosphate reductase 1 (GMPR1). The crystal structure data of GMPR1 was from protein data bank (PDB ID: 2BLE) with resolution of 1.9 Å. Structures of 1174 drugs were retrieved from database DrugBank. The docking was implemented with AutoDock Vina. GMPR1 and drug are used as receptor and ligand, respectively. The grid box accommodates all the atoms of GMPR1. The search space volume is 64×42×72 Å. The optimal drug was chosen by sorting the docking affinity reported by AutoDock Vina. The optimal drug has a more negative free energy (∆G). 3D structure of each drug was generated with Babel. **A-E**. Shown are the docked complex structures of the five ligands and GMPR1; A(b), Lumacaftor; B, Eltrombopag; C, Ledipasvir; D, Lurasidone; E, Deslanoside. In each panel, the left shows the full view of the complex and the right indicates the interaction details including the contacts between the GMPR1 and the ligands. H-bond is indicated as a green stick. ∆G means the docking affinity. P-value is estimated from the distribution of the docking affinities for 1174 ligands.

### Therapeutic effect of lumacaftor to AD

To this end, we tested the therapeutic effect on AD model mouse. The levels of β-Amyloid and phosphorylated Tau were determined with immunohistochemistry on 0 day, 10 days and 20 days after oral administration of lumacaftor. In hippocampus of the control mice, area of β-Amyloid (brown spot) increases with time (left panels of Fig. 5A). In the treatment mouse, the area does not increase so much as that in control in 20 days (right panels of Fig. 5A). Phosphorylated Tau (brown + blue spot) is drastically reduced after treatment in 20 days in this case (Fig. 5B). We estimated average area of β-Amyloid and counted average number of phosphorylated Tau-positive (PHF-1^(+)^) nerve cells in parietal lobe, temporal lobe and hippocampus (Fig. 5C-D and Tab. S4-5). The results indicated that the accumulation of β-Amyloid was greatly slowed down and phosphorylation of Tau was almost cleared, suggesting a satisfying effect.

**Figure 5.**
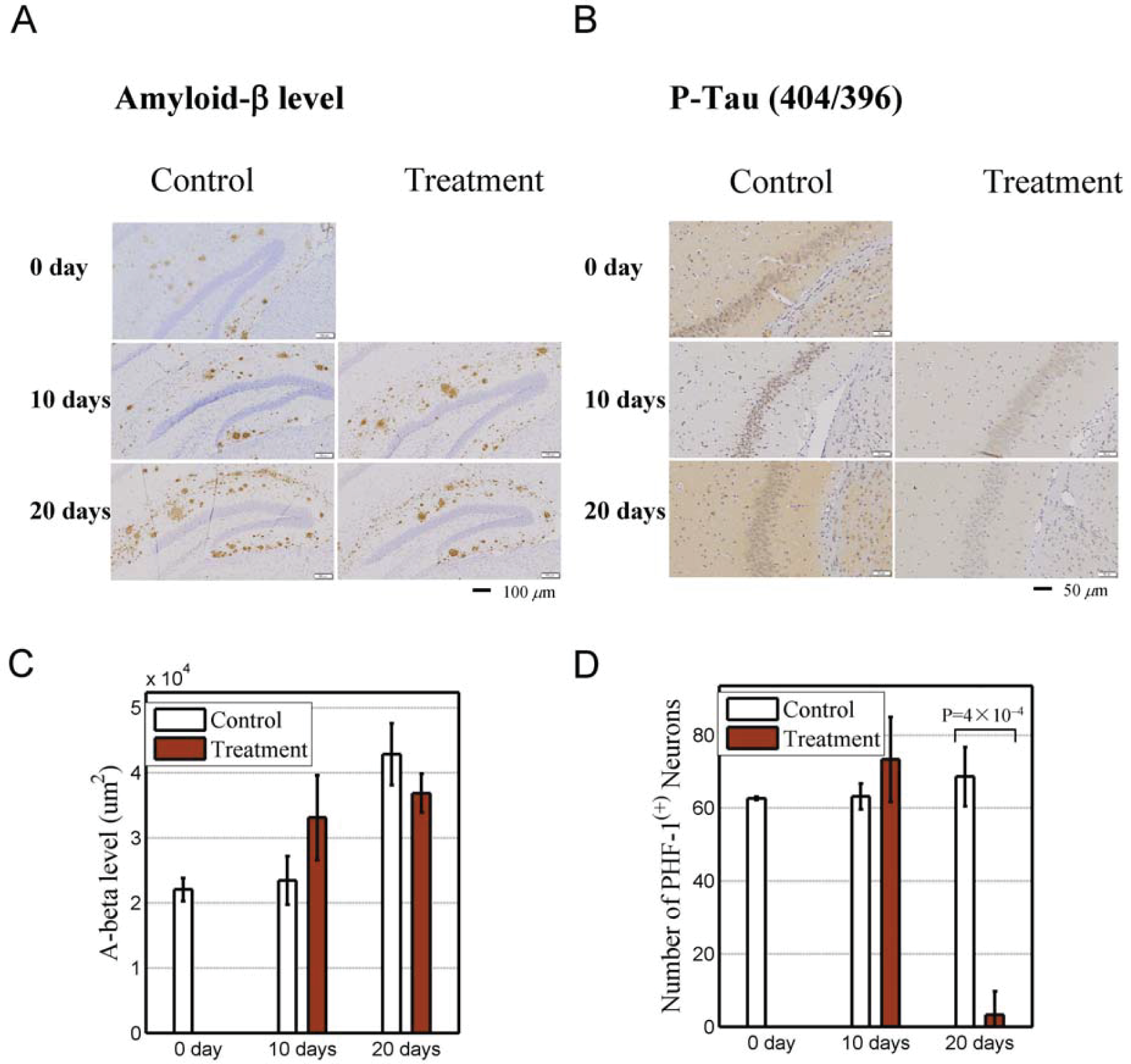
Therapeutic effect of lumacaftor on AD mice. Control group, six AD mice without lumacaftor in food. Brain tissues of any two mice were sectioned on the first day, the 10th day and the 20 day since the beginning of oral administration of the drug, respectively. Eight mice (treatment group) that were treated with lumacaftor were sectioned on the 10th day and the 20 day day. Four mice were executed each time. Levels of β-Amyloid and phosphorylated Tau represent with the averaged levels in three blindly-selected zones in parietal lobe, temporal lobe and hippocampus, respectively. **A**. Immunohistochemistry of β-Amyloid (brown spots) in hippocampus in control (left panels) and treatment mice (right panels). **B**. Same with subplot A except for phosphorylated Tau (brown + blue spots). **C**. Average area of β-Amyloid in parietal lobe, temporal lobe and hippocampus. Error bars indicate standard deviation. **D**. Average number of phosphorylated Tau-positive (PHF-1^(+)^) nerve cells in parietal lobe, temporal lobe and hippocampus. Error bars indicate standard deviation.

Taken together, this evidences that GMPR1 can be therapeutic target and lumacaftor does have therapeutic effect on AD, especially in preventing accumulation of β-Amyloid and in eliminating tau phosphorylation.

## Discussion

Molecule pathological mechanism is crucial to finding effective therapeutic strategy to AD. In this work, we identified some new biomarkers and therapeutic targets by analyzing gene expression difference in AD. Targeting one of the targets, we screened the potential inhibitors and tested the therapeutic effect in AD mice. Four points should be highlighted. The first is that we identified the genes with a different expression in AD case rather than in old population (≥ 60 yr). The expression changes of the genes represent more essential alteration in AD. After all, not all the old people exhibit AD symptom. Among the genes, GMPR is special because its expression increases in AD, which allows it to be a biomarker. The logistic model based on GMPR shows a good capacity of classifying AD cases (AUC > 60% in either dataset). The second is that the genes associate multiple pathways. Signaling of MET and WIF1 converges on GSK3β through different paths [3, 7]. But WIF1 seems to have a protective effect since its expression is down regulated, which leads an enhancement of Wnt signaling and a less phosphorylation level [5]. GMPR1 links with multiply pathways including AMPK and adenosine (A1/A2) receptor pathways. The activation of the pathways results in phosphorylation process [14, 29]. An elevated GMPR1 level means a higher phosphorylation level of Tau. Additionally, we inferred an abnormal transcriptional regulation at one JUND-binding enhancer of GMPR in AD. All of the above indicates GMPR1 is key node in molecule network of AD. The third is that in docking with GMPR1, lumacaftor, a FDA-approved drug, exhibits a high affinity and has good properties, and thus is a good candidate of inhibiting GMPR1. The last is that we tested the therapeutic effect of lumacaftor in AD model mice. The result showed that the drug can efficiently reduce the phosphorylation of Tau in both lobe and hippocampus.

## Datasets and method

### Datasets

Three sets of expression data were used in the analysis. The first dataset, which is microarray expression data including 32 AD cases and 47 non-AD cases from the postmortem human brain tissues of AD, was retrieved from literatures (Gene Expression Omnibus (GEO) accession ID: GSE36980) [18]. This data was used to identify AD-specific gene expression alteration. The second dataset is expression data including 22 old (≥ 60 yr, µ = 86, σ =12) and 19 young (<60 yr, μ = 35, σ =9.5) cases of the postmortem neuropathologically normal brain tissues from the frontal cortical regions. It was retrieved from literature (GEO ID: GDS5204)[23]. This dataset is used to identify the differential expression genes in old population. The third dataset is for 8 non-AD and 22 AD cases, which was retrieved from literature (GEO ID: GSE28146)[24]. The 22 AD cases consist of 7 incipient cases, 8 moderate cases and 7 severe cases. The dataset is used to validate the expression for the differential expression genes that was found in the first and third datasets.

The crystal structure datasets of GMPR1 was from protein data bank (PDB) (accession ID: 2BLE). The resolution is 1.9 Å. Structures of 1174 FDA-approved drugs were retrieved from database DrugBank [25].

### Differential expression analysis

Two-sample *t*-test was used to respectively detect the up- and down-regulated genes in two datasets, namely dataset GSE36980 (32 AD vs. 47 non-AD cases) and dataset GDS5204 (22 old people vs. 19 young people cases). The differentially expressed genes were identified using the criteria that p-value of *t*-test is ≤ 10^−5^ and the absolute value of log2 (fold change) is ≥ 0.1 for dataset 1 and is ≥ 0.15 for dataset 1. By comparing two sets of the differentially expressed genes, we identified the age-independent differentially expressed genes in AD. Volcano plot was employed to represent both the p-value and the fold change.

Logistic regression models were constructed with the differentially expressed genes to further evaluate the difference of the genes expression in datasets GSE36980 and GSE28146. The receiver operator characteristic (ROC) curves was used to show the performance of the models.

Co-expression genes were identified by calculating Pearson correlation coefficient (r) of the expression between a specific gene, for instance GMPR, and each of other genes in the first dataset. An enrichment analysis for both top 100 co-expression genes with gene GMPR and top 100 anti-expression genes with GMPR was carried out on both the terms of gene ontology (GO) and the pathways of KEGG with bioinformatics tool DAVID [26].

### Screening of inhibitors for GMPR

The inhibitors were screened for GMPR1 (PDB ID 2BLE) from 1174 FDA-approved drugs. The screening was carried out with molecule docking tool AutoDock Vina [27]. 3D structure of each drug molecule was generated with Babel (version 2.4.0) [28]. In docking, GMPR1 and drug molecule are used as receptor and ligand, respectively. The grid box accommodates all the atoms of GMPR1. The search space volume is 64×42×72 Å. The optimal drug was chosen by sorting the docking affinity reported by AutoDock Vina. The optimal drug has a more negative free energy (∆G). Empirical P-value is estimated from the distribution of the docking affinities for 1174 ligands.

### Immunohistochemistry of β-Amyloid and phosphorylated tau

Fourteen 10-month mice of B6C3-Tg (APPswePSEN1dE9Nju)/Nju were from Nanjing Biomedical Research Institute of Nanjing University (Nanjing, China). Inhibitor lumacaftor (VX-809) was purchased from Selleck Chemicals (Texas USA). Antibody β-Amyloid (D12B2) Rabbit mAb #9888 was from CST Biological Reagents company (Shanghai, China). Antibody of Tau phosphorylated sites 404/396 (anti-PHF-1 antibody, ab184951) was from Abcam (Shanghai, China).

Eight mice, test group, were given 0.3332 mg of lumacaftor (30% PEG400 + 0.5% Tween80 + 5% Propylene glycol + 64.5% double distilled water) with oil-containing food twice a day. Six mice, control group, were fed with same food except adding lumacaftor. On day 0, 10 and 20 after the administration with lumacaftor, the levels of both beta-amyloid protein and phosphorylated tau were tested with immunohistochemistry. Level of Aβ was represented with the average positive area of Aβ in three blindly selected zones in parietal lobe, temporal lobe and hippocampus, respectively under 10X field of view. Similarly, the number of the neurons with phosphorylated Tau was counted in three blindly selected zones (20X field of view) in parietal lobe, temporal lobe and hippocampus, respectively, and then averaged.

## Acknowledgments

This work was supported by the National Natural Science Foundation of China (No.31371339 and No.81660471).

## Authors’ contributions

Both authors contributed equally to the work. Both authors read and edit the final manuscript.

## Supplementary materials

### Tables

**Table S1.**
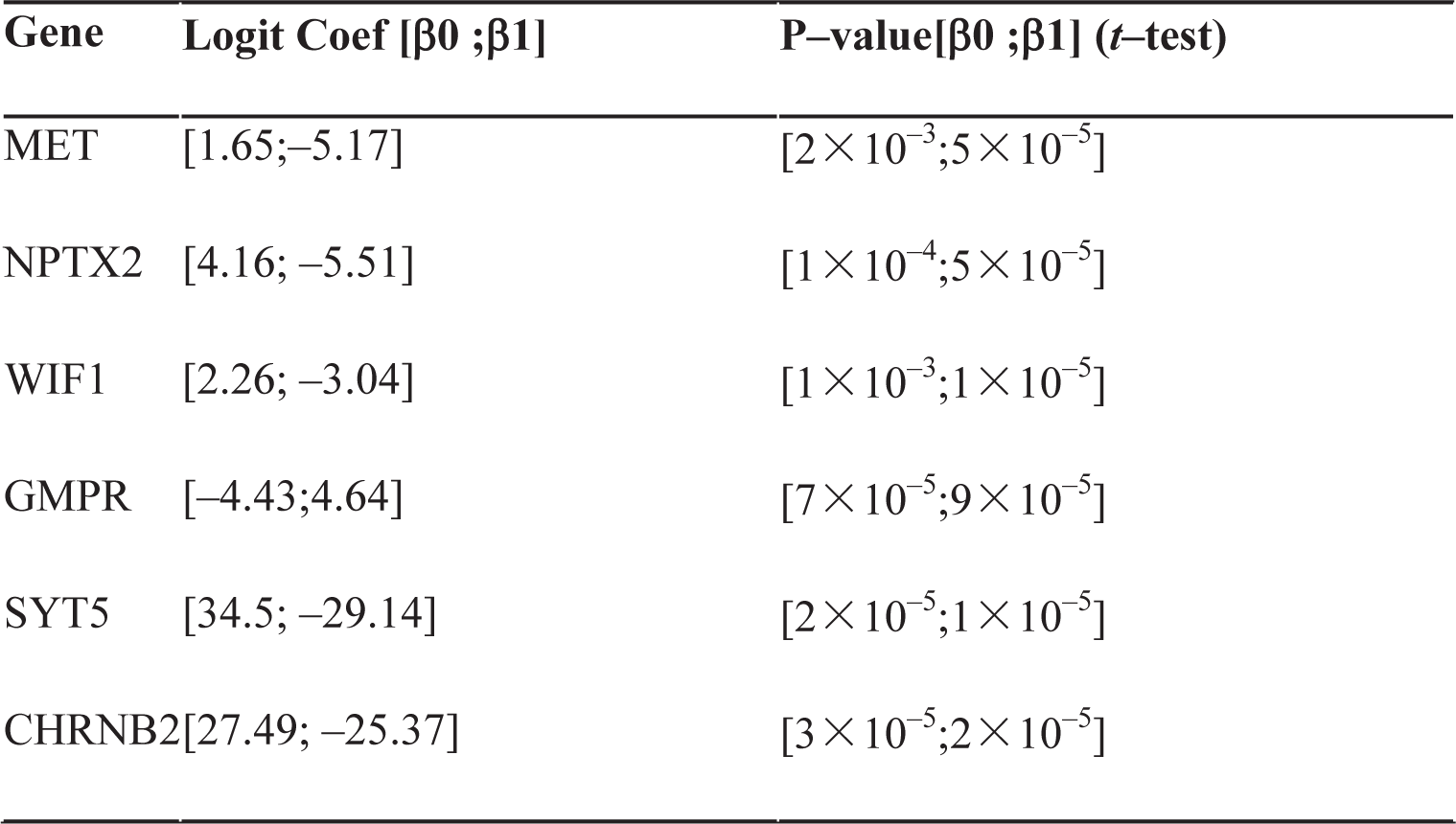
Logistic regression coefficients of each single gene-based model; P-value is the hypothesis test for the coefficients

**Table S2.** Logistic regression coefficients of the models using the genes combination; P-value is the hypothesis test for the coefficients.

| **Genes** | **Logit Coef [$\beta$]** | **P-value [$\beta$] (*t*-test)** |
| --- | --- | --- |
| [GMPR; WIF1] | [6.98;12.33;-23.34] | [0.2;0.02;0.007] |
| [NPTX2;GMPR] | [-2.12;21.44;-16.69] | [0.8;0.02;0.06] |
| [MET;WIF1;SYT5] | [-82.92;7.96;18.43;104.20] | [0.001;0.4;0.02;0.01] |
| [MET;NPTX2;WIF1;SYT5] | [-83.31;6.77;7.60;10.49;106.13] | [0.0008;0.5;0.4;0.1;0.007] |
| [MET;NPTX2;GMPR;SYT5] | [-61.53;6.43;11.51;-12.87;90.90] | [0.02;0.6;0.3;0.2;0.02] |
| [MET;NPTX2;WIF1;GMPR;SYT5] | [-132.30;17.71;2.02;24.11;-7.25;107.19;66.47] | [0.003;0.2;0.8;0.0;0.7;0.09;0.2] |

**Table S3.** Drugs with a good affinity with GMPR1.

| **Drug** | **Affinity**<br>**(Kcal/mol)** | **P-value**<br>**(n=1174)** | **Molecular formula** | **Molecular weight** |
| --- | --- | --- | --- | --- |
| **Ledipasvir** | **-9.10E+00** | **3.00E-03** | **C49H54F2N8O6** | **888.4134379** |
| Elbasvir | -8.80E+00 | 4.50E-03 | C49H55N9O7 | 881.4224451 |
| Venetoclax | -8.50E+00 | 8.70E-03 | C45H50ClN7O7S | 867.3180959 |
| Pasireotide | -8.50E+00 | 8.70E-03 | C58H66N10O9 | 1046.501424 |
| **Deslanoside** | **-8.50E+00** | **8.70E-03** | **C47H74O19** | **942.4824302** |
| **Eltrombopag** | **-8.40E+00** | **8.70E-03** | **C25H22N4O4** | **442.1641052** |
| Suramin | -8.4 | 8.70E-03 | C51H40N6O23S6 | 1296.046906 |
| Posaconazole | -8.3 | 0.0129 | C37H42F2N8O4 | 700.3297083 |
| Ponatinib | -8.20E+00 | 1.29E-02 | C29H27F3N6O | 532.2198441 |
| Plicamycin | -8.20E+00 | 1.29E-02 | C52H76O24 | 1084.472653 |
| Simeprevir | -8.20E+00 | 1.29E-02 | C38H47N5O7S2 | 749.2916903 |
| Lifitegrast | -8 | 0.0223 | C29H24Cl2N2O7S | 614.0681277 |
| Ombitasvir | -8 | 0.0223 | C50H67N7O8 | 893.5051121 |
| **Lumacaftor** | **-8** | **0.0223** | **C24H18F2N2O5** | **452.118378** |
| Daclatasvir | -8 | 0.0223 | C40H50N8O6 | 738.3853314 |
| Regorafenib | -7.90E+00 | 2.23E-02 | C21H15ClF4N4O3 | 482.0768809 |
| **Lurasidone** | **-7.90E+00** | **2.23E-02** | **C28H36N4O2S** | **492.2558971** |
| Nilotinib | -7.90E+00 | 2.23E-02 | C28H22F3N7O | 529.183793 |

**Table S4.** Average area of β-Amyloid (+) in each AD mice (μm^2^).

| Data |  | Control |  | Treatment |  |  |  |
| --- | --- | --- | --- | --- | --- | --- | --- |
| July 19, 2017 | Animal ID | 1943 | 1641 | / | / | / | / |
| | Aβ (+) Area | 20759.369 | 23273.615 | / | / | / | / |
| July 29, 2017 | Animal ID | 1950 | 1856 | 1639 | 75 | 19 | 52 |
| | Aβ (+) Area | 26054.681 | 20793.338 | 34039.558 | 39110.784 | 35329.279 | 23860.400 |
| August 8, 2017 | Animal ID | 1951 | 1931 | 1836 | 21 | 50 | 24 |
| | Aβ (+) Area | 39475.639 | 46210.828 | 35653.080 | 33247.865 | 39672.366 | 38833.005 |

**Table S5.** Count of PHF-1^(+)^ neurons in each AD mice.

| **Data** |  | **Control** |  | **Treatment** |  |  |  |
| --- | --- | --- | --- | --- | --- | --- | --- |
| July 19,<br>2017 | Animal ID | 1943 | 1641 | / | / | / | / |
|  | Count of PHF-1<sup>(+)</sup><br>neurons | 62.89 | 62.33 | / | / | / | / |
| July 29,<br>2017 | Animal ID | 1950 | 1856 | 1639 | 75 | 19 | 52 |
|  | Count of PHF-1<sup>(+)</sup><br>neurons | 60.67 | 65.67 | 73.67 | 88.33 | 60.00 | 71.33 |
| August 8,<br>2017 | Animal ID | 1951 | 1931 | 1836 | 21 | 50 | 24 |
|  | Count of PHF-1<sup>(+)</sup><br>neurons | 74.33 | 62.89 | 0.00 | 0.00 | 0.00 | 13.00 |

### Supplementary Figures

#### Figure legends

**Figure S1.**
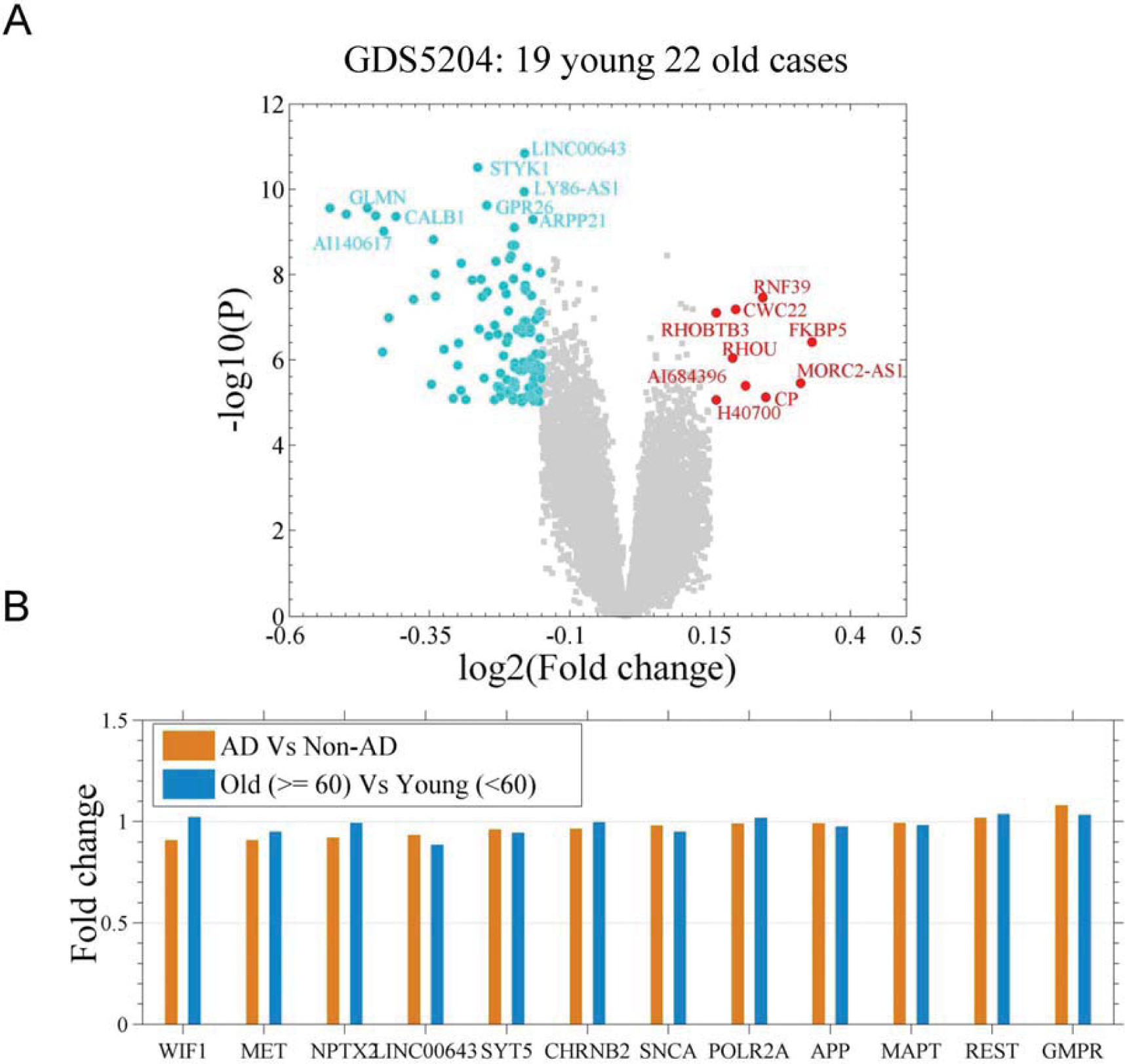
**A** differential expression analysis between young and old population. A. A differential expression analysis between 22 old (≥ 60 yr, μ = 86, σ =12) and 19 young (<60 yr, μ = 35, σ =9.5) samples of the postmortem neuropathologically normal brain tissues from the frontal cortical regions (GDS5204); shown are fold change (log_2_) against p-value (log10) (two-sample t-test). The cutoff for the differentially expressed genes is p-value ≤ 10^−5^ and either log2 (fold change) ≥ 0.15 or ≤ −0.15. **B.** The fold change degree of the genes REST, MAPT, APP, SNCA, POLR2A, GMPR, WIF1, NPTX2, MET, LINC00643, SYT5 and CHRNB2 in two comparisons (AD vs non-AD and old vs. young).

**Figure S2.**
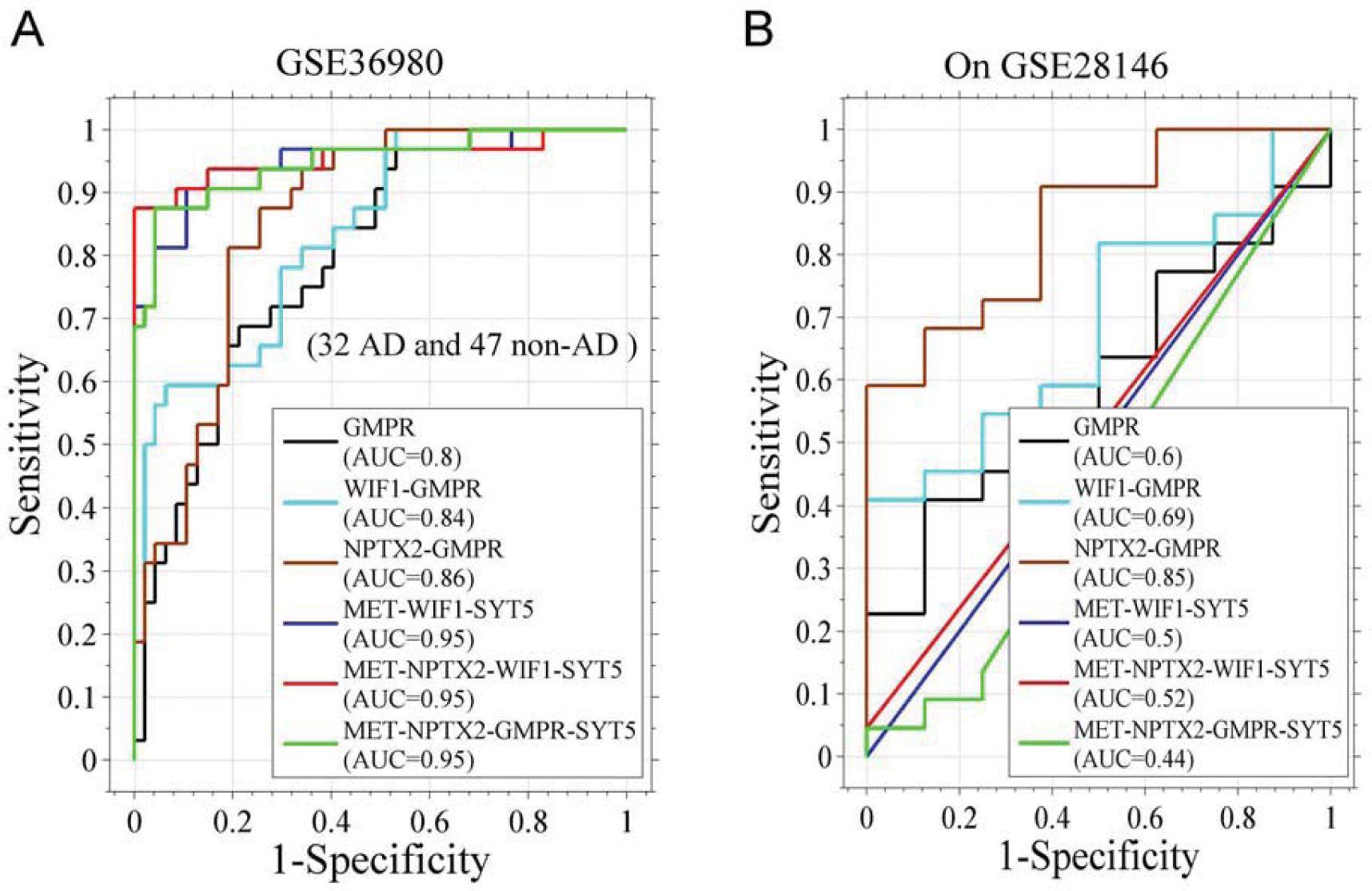
Performance of the logistic regression models based on the combination of the genes that are different in expression in AD. The regression parameters are listed in Table S2. **A.** The receiver operator characteristic (ROC) curves of the models on dataset of GSE36980 (32 AD and 47 non-AD samples). AUC means area under the curve. **B.** ROCs of the models on dataset GSE28146 (8 non-AD and 22 AD samples).

**Figure S3.**
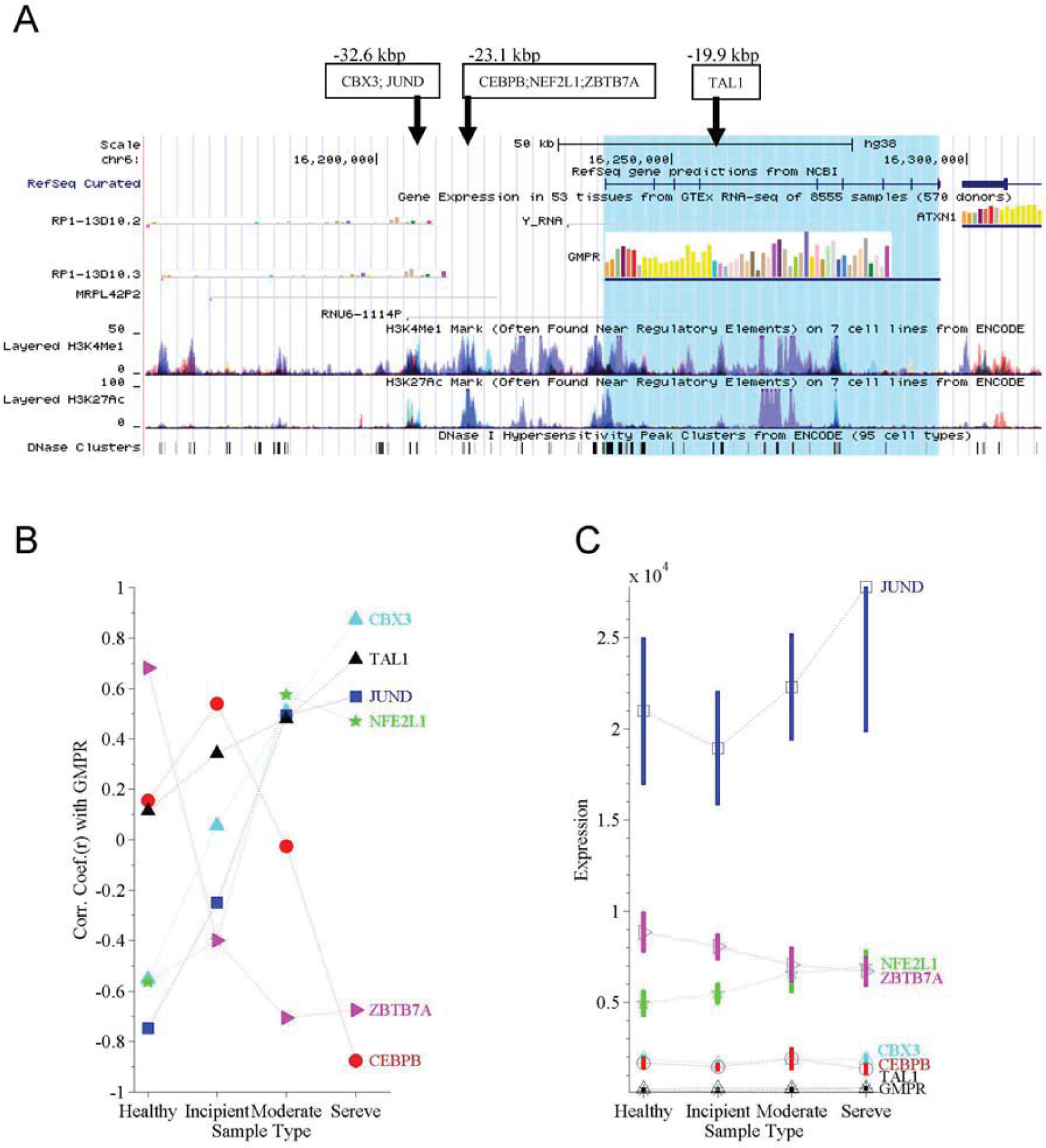
Transcription factor JUND regulates GMPR through binding at an enhancer. **A.** Shown are enhancers and transcription factors (TFs) around gene GMPR. Enhancers are indicated with black arrows. They have both acetylation of lysine 27 on histone H3 (H3K27ac) and mono-methylation of lysine 4 on histone H3 (H3K4me1). **B.** Pearson correlation coefficients (r) of expression between GMPR and the gene encoding TFs. The correlation coefficients are calculated in 8 healthy cases, 7 incipient AD cases, 8 moderate AD cases and 7 severe AD cases, respectively (GSE28146). The correlation coefficient for JUND changes from a negative value in healthy cases to a positive value in severe AD cases. C. Expression levels of genes GMPR, CBX3, JUND, ATL1, ZBTB7A, NF2L1 and CEBPB. Vertical bar indicates standard deviation error.

**Figure S4.**
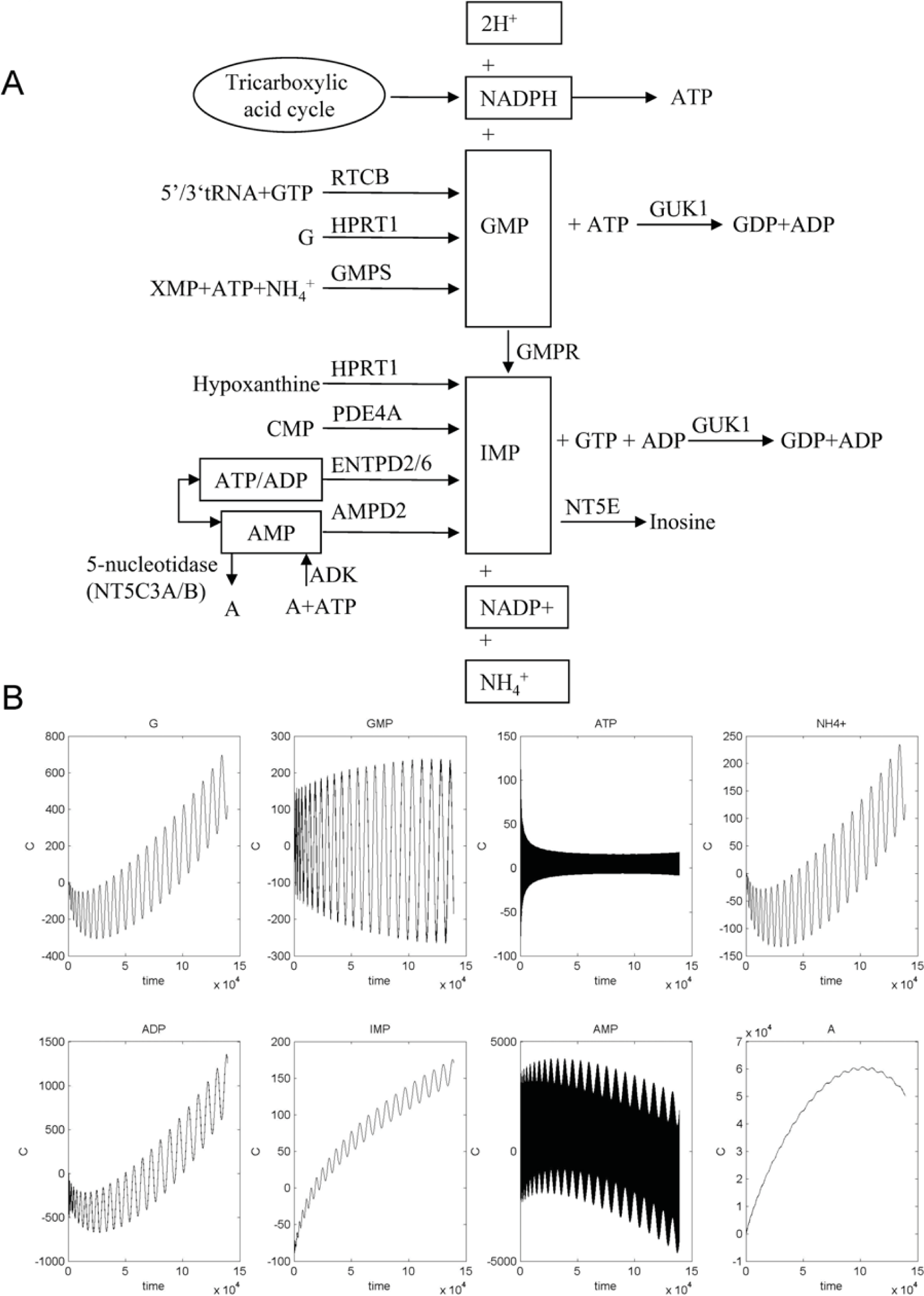
GMPR1is responsible for the reaction of GMP to IMP. Adenosine (A), AMP, ATP, ADP, GTP and other molecules connect the reaction. **A.** Reactions that link with either the reactants or the resultants of the GMPR1’s reaction. **B.** A dynamics simulation for the network that is constituted with the reactions shown in A. For each reaction, a partial differential equation is used to represent the relationship between reactant and resultant. Reaction rate constant is assigned in according to gene expression data of the enzymes. We let concentration of ATP oscillate with time and observe the dynamics of other molecules.

**Figure S5.**
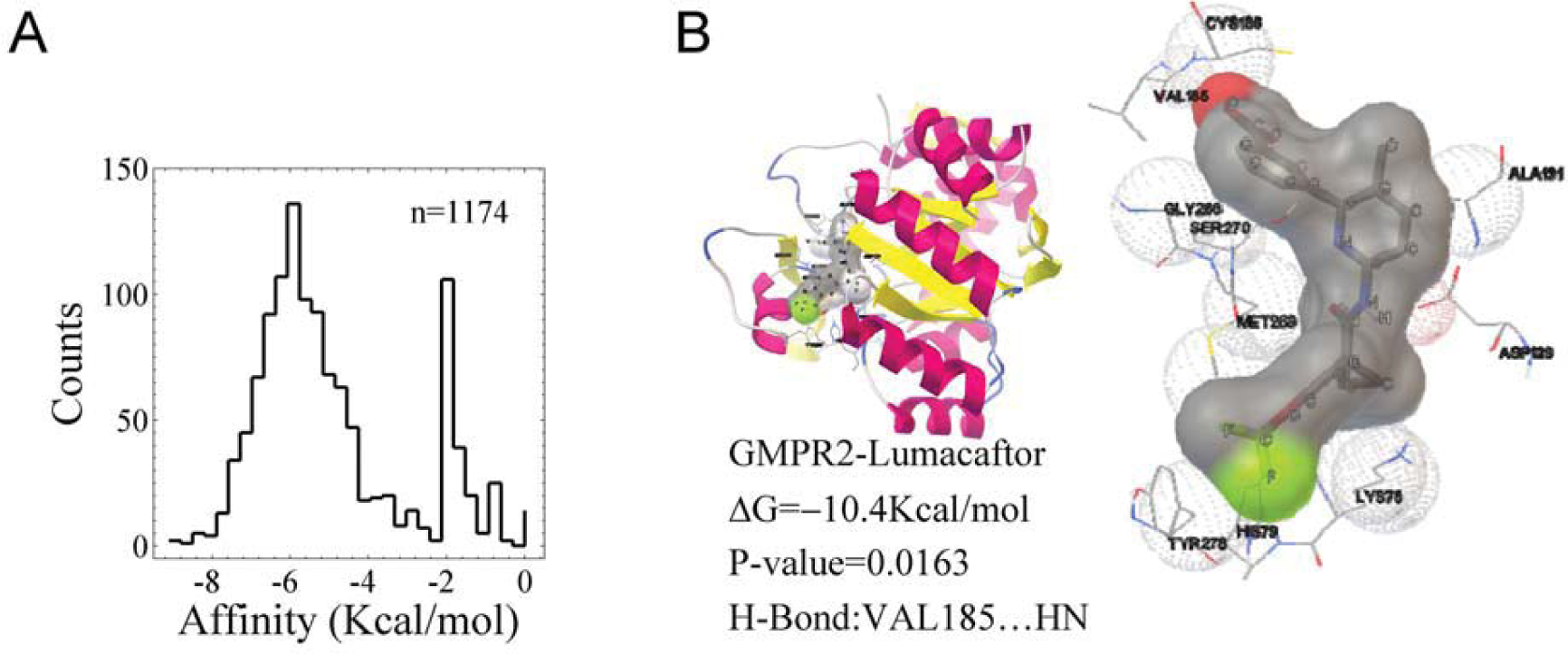
**A**, The affinity distribution of 1174 drugs in docking with GMPR1. For each drug, AutoDock Vina reports 9 conformations. The conformation with the most negative affinity was chosen and the affinity value was used to calculate the distribution. **B**, The docked complex structures between human guanosine monophosphate 2 reductase 2 (GMPR2) and Lumacaftor. The crystal structure data of GMPR2 is retrieved from protein data bank (PDB ID: 2C6Q, resolution = 1.7 Å). Other computation is similar as that for GMPR1.

